# Language may be all omics needs: Harmonizing multimodal data for omics understanding with CellHermes

**DOI:** 10.1101/2025.11.07.687322

**Authors:** Yicheng Gao, Weixu Wang, Yuheng Zhao, Kejing Dong, Caihua Shan, Weizhong Zheng, Till Richter, Zekai Li, Siming Chen, Fabian J. Theis, Qi Liu

## Abstract

Decoding cellular systems requires integrating diverse omics data, yet most models are trained from scratch on a single modality, restricting generalization. Here we present CellHermes, a biological language model that leverages pretrained large language models (LLMs) to integrate multimodal forms of omics data, such as transcriptomic profile and PPI network, through natural language for better omics understanding. By reformulating these datasets into question-answer pairs, CellHermes emulates multiple self-supervised learning paradigms within a single, universal, language-based framework with LORA fine-tuning on existing natural language-LLM, achieving comparable or even better performance compared to current single-cell foundation models. Within this framework, CellHermes functions as an encoder, a predictor and an explainer for supporting a range of downstream tasks. For encoder, CellHermes encodes biological entities with embeddings that enable accurate network discovery and cross-dataset generalization; For predictor, CellHermes unifies heterogeneous downstream tasks by translating them into natural language question-answers pairs, allowing a single model to perform multi-task prediction by instruction fine-tuning; For explainer, CellHermes explains molecular mechanisms by combining attention analysis with text-based reasoning, leveraging the interpretability and reasoning capabilities of LLMs. We evaluated the ability of CellHermes as an *encoder* on representing genes and cells using 5 gene-level downstream tasks and 5 diverse single-cell datasets across different tissues compared with other single cell foundation models. We also evaluated CellHermes as a *predictor* on BioUniBench, a new benchmark of 10 tasks across 7 databases for LLMs, including perturbation response, cell fitness estimation, gene-disease association, and cell type identification. All benchmarks suggest CellHermes can unify multiple biological tasks within one model without compromising performance. By applying CellHermes as an *explainer* for a melanoma patients dataset, it can uncover potential key genes on tumor-reactive T cells. Together, CellHermes establishes natural language as a unifying medium for omics, offering a foundation that, in the future, potentially enables a more integrated and interpretable research loop of biological representation, prediction, and interpretability.

## Introduction

In the rapidly evolving landscape of biological research, omics knowledges, including sequenced tabular datasets, from multi-omics and related fields, has accumulated into vast and complex datasets that capture multiple layers of biological regulation and function^1–4^; yet their large scale, heterogeneity and multi-modal nature pose great challenges for integrative analysis with traditional domain-specific models, limiting systematic understanding of regulatory mechanisms across multiple scales and layers of the system. Despite their diversity, the basic sequence structure of molecules like DNA and continuous biological processes can be conceptualized as a symbolic system, akin to a *biological language* of life^5,6^. In recent years, large language models (LLMs), originally developed for natural language processing, have exhibited strong capabilities in sequential modeling, including text representation^6–9^, natural language understanding and reasoning^10,11^. This convergence invites a compelling question: Can we ‘translate’ structural biological omics data into natural language, thereby enabling pretrained natural language-LLMs – with their embedded world knowledge and reasoning skills – to comprehend and analyze cellular systems?

Omics data is broadly categorizable into two forms, purely tabular entries, such as transcriptomic profile, and nodes connected by edges in a graph, organized as a biological interaction knowledge graph such as Protein-protein interaction (PPI) network. Modeling efforts have focused on both forms. For tabular data, initial approaches^12,13^ used variational autoencoders (VAEs) for cell representation or other generative models^14–16^ to model gene expression changes. For graph data, graph neural networks (GNNs) have been a primary tool applied to modeling molecular level or cell level interaction^17–19^. Recently, a significant trend is building foundation models for omics data. This includes adapting BERT or GPT architectures^5,20–25^ for single-cell transcriptomic data using masked language modeling (MLM) or autoregressive learning. This trend also encompasses employing graph-based supervised techniques to learn gene embeddings for graph-structured omics data^26^. While these strategies have shown considerable success, they are often constrained by their reliance on specialized, modality-specific model architecture. This can result in representations that are not easily integrated across different omics layers or with textual knowledge, potentially limiting their capacity for reasoning and generalizability.

Unlike traditional tabular– or graph-based methods^18,20,21,26^, natural language provides three key advantages for omics, potentially providing a fundamental path to unify the complex forms of omics. First, it offers a flexible tokenizer scheme. By representing diverse biological entities, such as gene expression or gene interactions, as discrete tokens, we can structure them into sequences analogous to ‘sentences’^27^, therefore, language can inherently encode context-dependence and prior knowledge, thereby enhancing model generalization^28–31^. Second, language can readily describe the objective of tasks, unifying different tasks within one model and potentially facilitating the knowledge transfer between different downstream tasks by prior knowledge^32,33^. Third, pretrained natural language-LLMs already embody powerful mechanisms for reasoning, enabling human-readable explanations of system behaviors^34,35^. Together, these properties suggest that natural language is a practical unifying medium for modeling the complex of omics.

To realize this vision, we propose CellHermes, a general-purpose biological language model that unifies diverse omics data by leveraging the full capability of natural language-pretrained LLMs. Building on prior work that first introduced gene tokenization and Transformer pretraining for single-cell data (Geneformer^20^) and subsequently incorporated natural language–pretrained LLMs into single-cell modeling (Cell2Sentence^7^ and C2S-Scale^36^), CellHermes extends this foundation along two key axes. First, it applies parameter-efficient LoRA adaptation instead of full fine-tuning. Second, it broadens the data formulation to encompass graph-based and tabular omics within a multitask question-answering framework. In doing so, CellHermes harmonizes multimodal omics modalities, tabular, graph-based, and textual, to advance integrative omics understanding, grounded in the insight that diverse omics layers can all be expressed in language-like formats and aligned within a shared representational space of LLMs. CellHermes is first constructed by transforming multiple omics modalities, including transcriptomic matrices and gene interaction graphs, into natural language. Inspired by recent findings that natural language can emulate the abilities of GNNs^37^, we design instruction templates to encode self-supervised learning signals by question-answer pairs, including masked language modeling (as in BERT^38^) and autoregressive prediction (as in GPT^39^), alongside masked node and link prediction (as in GNNs)^40,41^. This novel formulation enables a single pretrained LLM to integrate heterogeneous omics data within one unified training framework, thereby enhancing its understanding of omics data from multiple views. The concept of representing single-cell transcriptomic profiles as sequences of ranked gene names and leveraging pretrained large language models for cell embeddings was first introduced in Cell2Sentence^7^, extending earlier ranked-gene representations from Geneformer^20^. CellHermes extends it toward a multimodal formulation that unifies both tabular (transcriptomic) and graph-structured (e.g., PPI) omics data within a question–answer–based multitask framework. Once constructed, CellHermes can then be applied across three functional modules: the embedding module (“encoder”), the downstream functional outcome module (“predictor”), and the natural language interpretation module (“explainer”). This design allows CellHermes to span from representation learning to prediction and interpretability. Across a series of benchmarks, we evaluate CellHermes’ modules. As an encoder, it generates high-dimensional embeddings of genes and cells, outperforming state-of-the-art single-cell foundation models, and supports the construction of cell-type-specific interaction networks for biological discovery. As a predictor, CellHermes leverages a natural language interface to integrate seven popular, heterogeneous databases across ten biological tasks, ranging from perturbation assays from Perturbase^42^ to cell fitness data from DepMap^43^, into a unified multitask objective for the construction of a universal cellular response predictor. Finally, as an explainer, CellHermes offers interpretability by combining attention-weight analysis within its architecture^44^ and text-based reasoning to highlight potential genes contributing to the cell behaviors.

## Results

### The CellHermes framework

Different from previous omics-specific foundation models, which are mostly built by pretraining a transformer architecture from a single omics modality^45^, CellHermes leverages existing natural language-LLM’ capacity and fine-tunes it for single-cell omics (Fig. 1a). Instead of training a new model, we unify omics data with natural language, translating biological resources into a question-answer format that LLMs can natively process and adapt to. With this framework, we bridge structured cell omics data, such as tabular data and graph structured data into the unstructured natural language domain, allowing a single LLM backbone to learn from and reason over multiple biological data types without a need for separate, modality-specific architectures. First, we designed a set of modality-specific instruction templates that translate structural biological data into textual descriptions that can be understood by LLM. Following Geneformer^6^ and Cell2Sentence^7^, a single-cell transcriptomic profile is represented as a gene-expression “sentence”^6,7^, where genes are tokens and quantitative measurements are transformed into gene name rank. Similarly, gene interaction networks are transformed into “relation statements”^37^ that capture edge connectivity (Fig. 1b). By unifying all modalities into a consistent question-answer format as illustrated in Fig. 1b, CellHermes allows a single LLM model to emulate multiple self-supervised learning paradigms simultaneously: (1) masked language modeling (MLM) for token-level feature masking, analogous to BERT pretraining; (2) autoregressive modeling for sequence generation, as in GPT; (3) masked node prediction for graph-structured omics data, and (4) masked link prediction for graph-structured omics data (Fig. 1b).

**Fig. 1.**
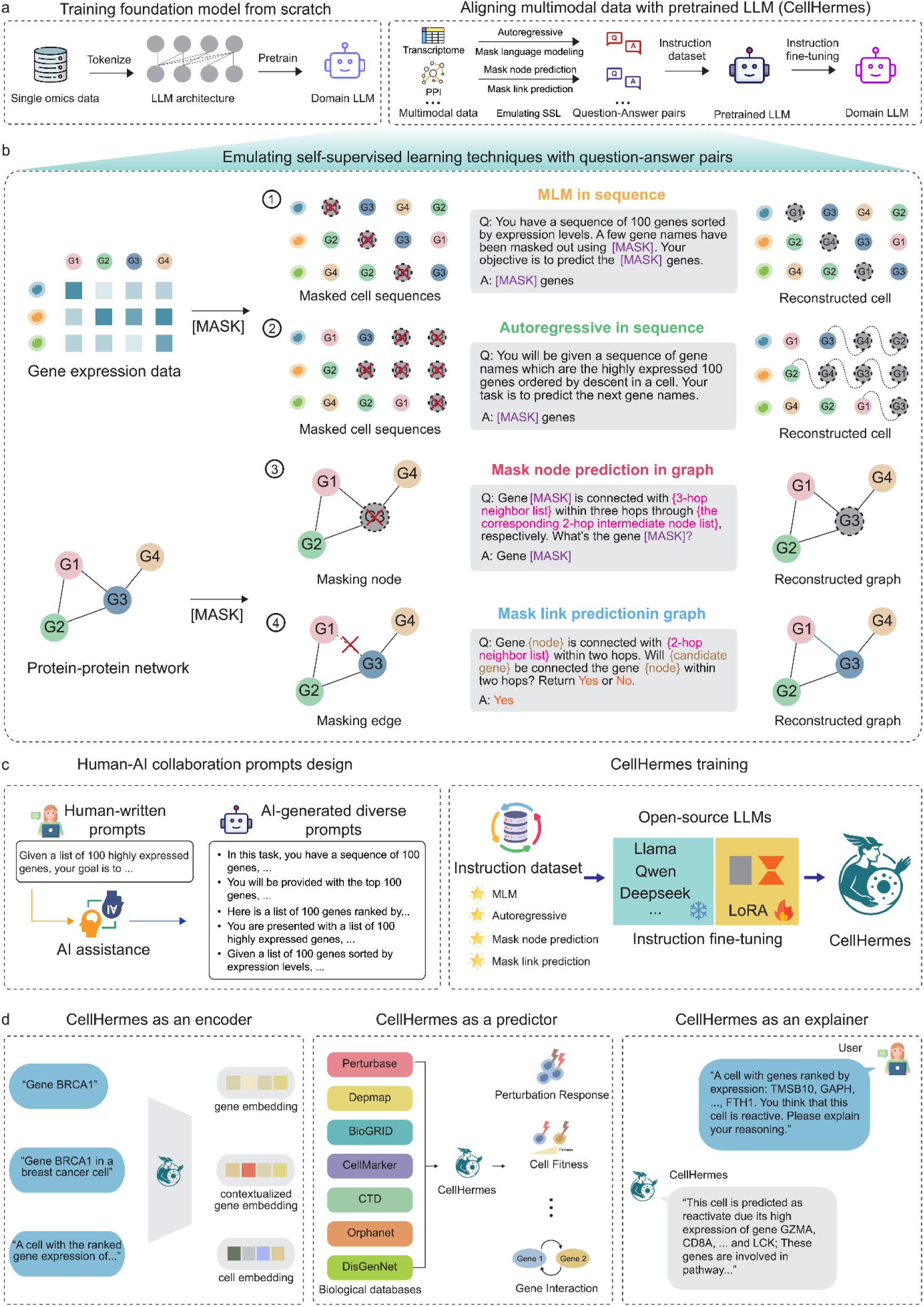
| CellHermes construction by integrating different self-supervised learning techniques. a. Comparative diagram of different foundation model construction between previous methods and our method; In most previous methods, we usually take one of the omics data and tokenize it for feeding into the pretraining model architecture (e.g. autoregressive-based GPT architecture), so that we can pretrain a omics-specific foundation model from scratch. In our method, we align multimodal omics data with pretrained LLM. First, we designed different question-answer pairs (instructions) for emulating the self-supervised learning techniques for each modality. Specifically, we emulate autoregressive and mask language modeling for transcriptomic data and emulate mask node prediction and mask link prediction for graph data (PPI network in this study). All of these instruction datasets then were together used for instruction fine-tuning the pretrained natural language-LLMs to adapt them into omic-specific LLM. b. Schematic illustration of emulating self-supervised learning strategies for tabular and graph-structured omics data by transforming omics data into a textual format and designing instruction templates for SSL tasks. In gene expression data, they are first transformed into sentence (ranking the gene names based on gene expression values) and masked language modeling (MLM) and autoregressive modeling are applied to represent genes and cells. In protein–protein interaction networks, the relations between genes are first transformed into natural language for constructing question-answer pairs. Then gene representations are learned through masked node prediction and masked link prediction. c. Construction of CellHermes. We begin with a human-written description, which then serves as an input to ChatGPT-4o for generating diverse instructions templates, capturing a broad range of natural human question-framing styles. Based on these constructed question-answer pairs from emulating self-supervised learning techniques (0.1 million instructions for transcriptome data and 0.2 million for PPI network in this study), we used low rank adaptation (LoRA) fine-tuning on the existing natural language-LLMs (we used LLaMA-3.1-8B-Instruct in this study). d. CellHermes can then be used as an encoder to encode genes and cells, as a predictor to perform multi-task prediction by integrating various biological databases, and as an explainer to perform reasoning based on its prediction results.

We selected transcriptomics data and protein-protein interaction (PPI) networks as representative examples of omics data for tabular and graph data, respectively, as they are widely available, biologically informative, and extensively benchmarked in prior studies^18,20,21,26^. We assembled a multimodal pretraining corpus, consisting of ∼1 million single-cell transcriptomics data across various tissues and cell types from CellxGene^4646,47^, and a protein-protein interaction network involving ∼ 20,000 genes^4^ (Supplementary Note 1). For each pretraining objective, we designed task-specific templates (Fig. 1b and Supplementary Note 1). To simulate masked language modeling (MLM), we designed two types of templates, differing in if the mask prediction depends on the meta information of the cell. For autoregressive modeling, we predict the next token from the preceding context. For masked gene prediction in graphs, we designed five template types based on varying hop-level neighborhood information. For masked link prediction in graphs, we developed ten template types, again varying by hop-level context (Methods and Supplementary Note 1). These diverse template types are mechanistically analogous to various GNNs, and as a result, they inherit their advantages^37^. To increase linguistic diversity and robustness, we used ChatGPT-4o^39^ to augment instruction types with 20 diverse task descriptions^48^ (Methods). This ensures that the model was exposed to varied phrasings for the same underlying biological query, improving generalization to unseen data. In total, we generated 2,272,436 question-answer pairs each for MLM and autoregressive tasks, 99,860 pairs for masked node prediction, and 1,455,846 pairs for masked link prediction (Methods and Supplementary Note 1). These datasets as a bunch of question-answer pairs, which include transcriptomic and PPI information, can then be jointly used to fine-tune or ‘pretrain’ existing LLMs with LoRA, using LLaMA-3.1-8B-Instruct^49^ as backbone in this study, resulting in a unified CellHermes model. (Fig. 1c and Methods). The number of parameters in CellHermes LoRA adapter is 20,971,520.

CellHermes readily offers an embedding model, enabling the representation of both genes and cells. Owing to the flexibility of natural language, contextual information, such as cell identity, can be seamlessly incorporated to generate context-aware embeddings. Beyond encoding, CellHermes functions as a predictor, capable of performing multi-task prediction within a single model without compromising performance. This is achieved by integrating diverse biological databases in text form and reformulating conventional supervised learning tasks into a question-answer framework. Finally, CellHermes can act as an explainer, leveraging its reasoning capabilities to provide interpretable explanations for its predictions (Fig. 1d).

### CellHermes as an encoder to embed biological entities

The design of CellHermes, which operate on text representations rather than modality-specific numerical format., offers a potential path to encode biological entities that can be expressed in natural language form, such as a gene, cell, pathway, perturbation, or interaction. This flexibility is enabled by the unification strategy that is intended to reformulate structured biological data into natural language, making it potentially interpretable by an LLM.In this study, we focus on gene and cell embeddings as illustrative examples, as they represent two of the most fundamental and widely studied biological entities. This makes them natural test cases for evaluating the capacity of CellHermes to capture biologically meaningful representations. To assess the potential for data leakage, we formulated evaluation tasks from the previous work^6^ as question-answer pairs and subjected them to direct, zero-shot inference by the LLM. The model’s inability to solve these tasks under these conditions indicates that data leakage was not a contributing factor to our results (Supplementary Note 2).

### Embedding genes and constructs into cell-specific gene networks

To encode genes, we used CellHermes as an encoder by taking the hidden embedding of the last token in the last transformer layers^9,50^ (Fig. 2a). Then, we evaluated the gene embedding through five downstream tasks, inspired by previous studies^6,20^, for predicting the role of genes: long versus short range TFs^20,51^, dosage sensitivity^52^, bivalent versus non-methylated genes^53,54^, Lys4-only-methylated versus non-methylated genes^53,54^ and gene-gene interactions^55^ (Fig. 2b and Supplementary Note 2). The performance was assessed by five-fold cross-validated ROC-AUC and PR-AUC with random forest (RF) or xgboost (XGB) classifiers^6^ using default parameters from scikit-learn package^56^. We benchmarked CellHermes against 9 existing single-cell foundation models, including scGPT^21^ and Cell2Sentence-27B^36^, as well as PCA^57^ and Random embeddings baselines (Supplementary Note 3). The results demonstrated that CellHermes consistently achieves superior performance with a dataset only consisting of ∼0.1 million single-cell transcriptome instructions and ∼0.2 million PPI network-based instructions (Fig. 2b and Supplementary Fig. 1). Notably, by using a small number of instructions to fine-tune existing natural-language LLM, CellHermes can outperform C2S-Scale^36^. Furthermore, we evaluated the effectiveness of the gene-ranking strategy for constructing cell sentences by comparing it with a random gene order baseline (Supplementary Fig. 2). We also examined the contribution of different data modalities, including transcriptomic profiles and the PPI network, and found that integrating both modalities yields a greater performance gain than using either alone (Supplementary Fig. 3). To assess the influence of data scale, we pretrained a series of CellHermes models using different amounts of transcriptomic dataset, from 0.05 million to the full dataset. We observed that even limited omics data were sufficient to yield high-quality pretrained embeddings while larger datasets may induce catastrophic forgetting in the underlying language model (Fig. 2c and Supplementary Fig. 3). This efficiency arises from two key factors. First, CellHermes leverages the generalization capacity of LLMs, thereby reducing the reliance on extremely large biological datasets. Second, by reformulating omics data into natural language, LLMs’ understanding of these biological entities can be enhanced by instruction fine-tuning, allowing meaningful representations.

**Fig. 2.**
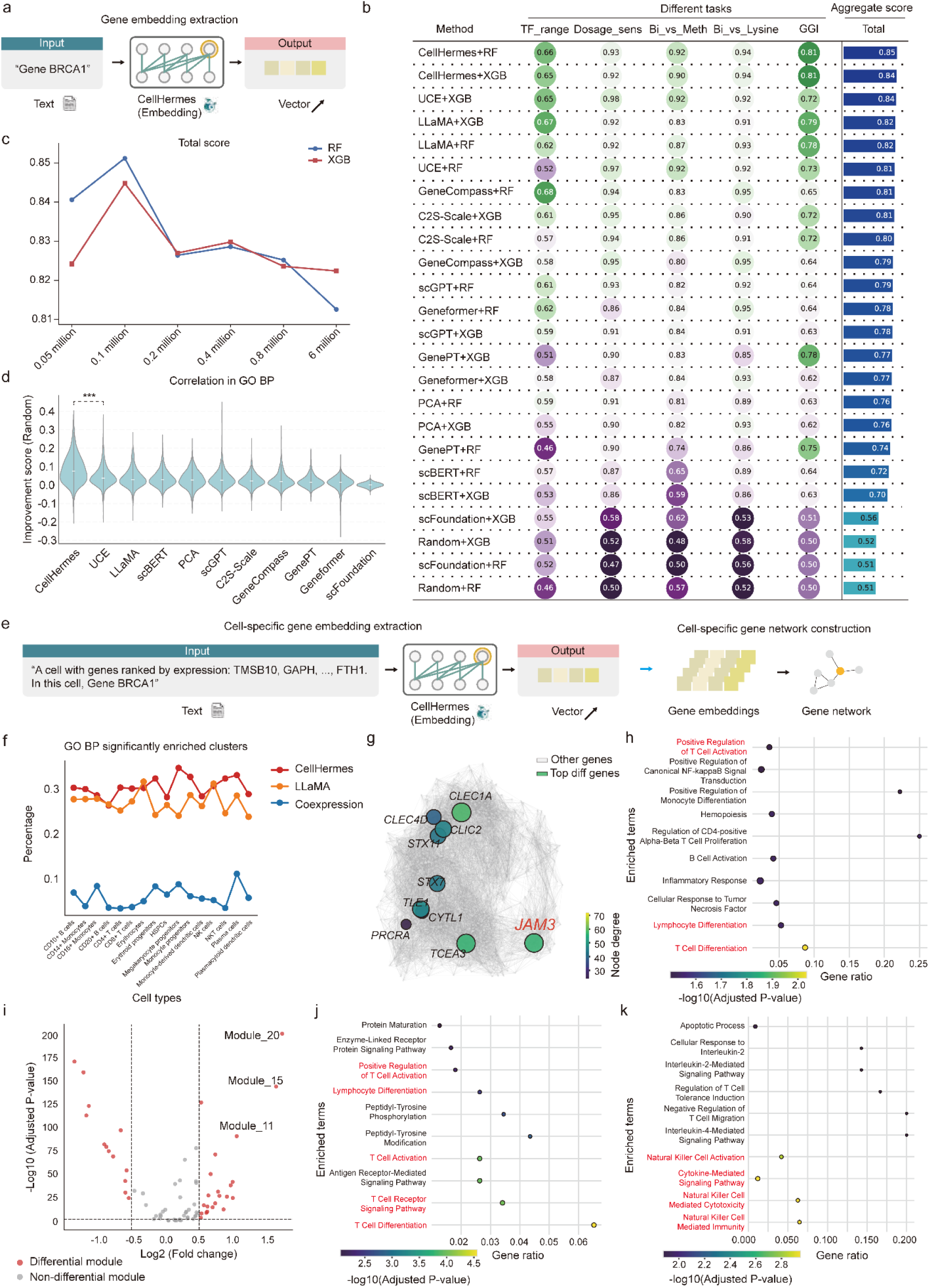
| CellHermes as an encoder for representing genes. a. Schematic illustration of transforming a generative model into an embedding model for representing genes by taking the last token embedding. b. ROC–AUC of benchmarked models on gene-level downstream tasks, including transcription factor range prediction (TF_range), dosage sensitivity prediction (Dosage_sens), bivalent chromatin vs. non-methylated genes (Bi_vs_Meth), bivalent chromatin vs. Lys4-methylated genes (Bi_vs_Lysine), and gene–gene interaction prediction (GGI). Aggregate scores were calculated as the mean across tasks. The performance of each model’s gene embedding was assessed by five-fold cross-validation with random forest (RF) or xgboost (XGB) classifiers using default parameters from scikit-learn package. c. Influence of pretraining dataset size on gene-level downstream tasks, quantified by meaning the ROCAUC scores achieved on each individual task. The compared models include the base LLaMA model without omics fine-tuning, a series of models trained on the instructions from PPI network together with increasingly large transcriptome datasets (0.05, 0.1, 0.2, 0.4, and 0.8 million, and the full dataset (6 million)), where the 0.1 million transcriptome model represents CellHermes in this study. d. Comparison of methods predicting Gene Ontology (GO) semantic similarity. Boxplots display the distribution of similarity scores (boxes show the 25th, 50th, and 75th percentiles). The “Improvement score (Random)” score is the performance gain over a random baseline, measured by the difference in Pearson correlation. e. Schematic illustration of incorporating cell information into instructions to obtain cell-specific gene embeddings, enabling the construction of cell-specific gene networks. This is a more concrete illustration for how we encode contextualized gene embeddings using CellHermes compared to Fig. 1d. f. Percentage of GO BP enriched clusters as a function of the cell types across different methods. g. Gene network of CD8⁺ T cells constructed from CD8⁺ T cell–specific embeddings, with highlighted genes showing the largest degree changes compared to the HSPC-specific network. h. GO BP enrichment analysis of the neighborhood genes of gene *JAM3*. i. Volcano plot of CD8+ T cell-specific gene modules. j. GO BP enrichment analysis of the genes in module 20. k. GO BP enrichment analysis of the genes in module 11.

Next, we evaluated the functional utility of the CellHermes gene embedding by adopting two approaches defined in the previous study^18^. First, to determine if the learned representations captured known functional relationships, we calculated the semantic similarity and co-embedding coefficient for every gene pair^18,58^. The semantic similarities were calculated based on different annotation databases, including GO Biological Process (GO BP)^59^, GO Cellular Component (GO CC)^59^, GO Molecular Function (GO MF)^59^, and KEGG pathway^60^. CellHermes gene embedding exhibited higher correlation compared to the existing single-cell foundation models (Fig. 2d, Supplementary Fig. 4 and Methods). Second, we evaluated the ability of embeddings to recover functional groupings by performing K-means clustering with cluster numbers ranging from 20 to 80^18^ (Methods). For each clustering configuration, we quantified the proportion of clusters significantly enriched for one or more terms, as determined by gene set enrichment analysis. Across all clustering scales, CellHermes achieved higher enrichment ratios than competing models, indicating that its embedding more effectively organized genes into functionally coherent groups (Supplementary Figs. 5a-d).

In addition to evaluating gene embeddings in a general context, we also examined cell-type-specific gene embeddings (Methods). Leveraging the flexibility of LLMs, we incorporated the transcriptomic information of one cell into the prompt, enabling the model to generate gene embeddings tailored to that cell (Fig. 2e). Using a human immune dataset^21,61^ as an example, we assessed the ability of these embeddings to recover functional groupings for each cell type. Across all cell types, CellHermes achieved higher enrichment ratios than competing models across multiple annotation databases (Fig. 2f and Supplementary Figs. 5e-g). Then, we explored the biological insights that could be gained by differential network analysis. We constructed a cell-type-specific network from gene embeddings using k-nearest neighbor (kNN) methods and compared the network topology between hematopoietic stem and progenitor cells (HSPCs) and CD8+ T cells. We identified the top 10 genes that with the largest changes in node degree between the two cell types (Fig. 2g). Notably, JAM3 exhibited a high degree in CD8+ T cells but a marked decrease in HSPCs. GO Biological Process enrichment analysis of its neighboring genes revealed that JAM3 is associated with T cell differentiation and activation (Fig. 2h).

We further investigated if the constructed networks could be used to detect cell-type-specific gene modules. For each cell type, we applied Louvain clustering^62^ to the cell type-specific networks to identify gene modules. We then calculated the Jaccard Index for every pairwise comparison of modules between HSPCs and CD8+ T cells, enabling the identification of cell type-specific modules (Supplementary Fig. 6 and Methods). In CD8+ T cell, we focused on the top three upregulated modules (Fig. 2i and Supplementary Fig. 7a), and GO Biological Process enrichment analysis indicated distinct functional specializations: for example, module_20 was enriched for T cell activation and differentiation (Fig. 2j), while module_11 was enriched for cytotoxic T cell mediated killing (Fig. 2k). Notably, when we repeated the same analysis pipeline using conventional co-expression networks (Supplementary Fig. 7b), no biologically meaningful modules were detected (Supplementary Fig. 7c and d), highlighting the advantage of CellHermes-derived networks for functional module discovery.

### Embedding cells while preserving biological signals and mitigating batch effects

To encode cells, we used the generative CellHermes model as an encoder by extracting the final token embedding from the last transformer layer, using as input a “cell sentence” consisting of ranked gene names (CellHermes-s) (Fig. 3a, Methods). We also evaluated an alternative representation in which cell embeddings were generated by taking the weighted average of gene embeddings according to their expression levels (CellHermes-w) (Methods). We compared CellHermes against ten existing single-cell foundation models^5,20–25,63,64^, two traditional single-cell embedding methods^12,65^, and two text-based representation method^6,49^ across five datasets: the Immune^61^, Aorta^66^, Lung^61^, Pancreas^61^, and PBMC^67^ datasets (Supplementary Note 4). Using the scib-metrics^61^, we quantified both biological signal preservation and batch effect removal. As expected, Scimilarity, trained on a 23.4-million-cell atlas, achieved the highest overall scores across all datasets (Fig. 3b and c). Notably, despite being trained on only 0.1 million single-cells, CellHermes-s consistently achieved top-four performance across all datasets, demonstrating its strong efficiency and competitive representation quality relative to large-scale foundation models (Fig. 3b, c and Supplementary Figs. 8-11). This also highlights the potential of using LLM as a medium to understand omics data.

**Fig. 3.**
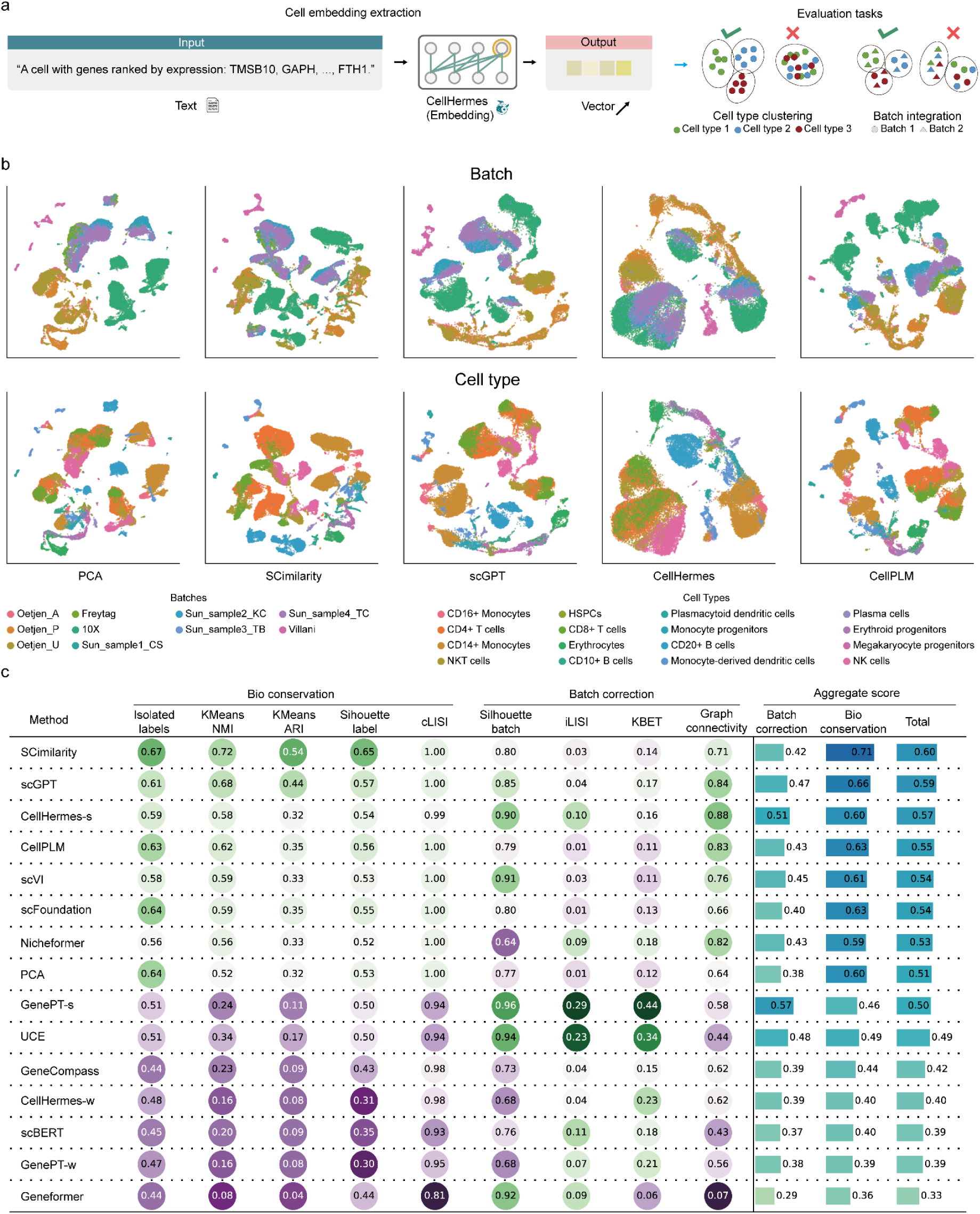
| CellHermes as an encoder for representing cells. a. Schematic illustration of transforming a “cell sentence” into a gene embedding with CellHermes, enabling cell type clustering and batch effect removal. This is a more concrete illustration for how we encode cell embeddings using CellHermes compared to Fig. 1d. b. UMAP visualization of different embedding methods, including four best-performing methods and one traditional PCA method, colored by annotated batch and cell type information. c. scIB integration scores for each benchmarking method on the Immune dataset, calculated by the *scib-metric* package; The last three columns indicate the average scores across all metrics. CellHermes-s and CellHermes-w embeddings were constructed following the protocol of GenePT. CellHermes-s uses the direct text embedding of a cell’s natural language description (“cell sentence”), while CellHermes-w is computed as the expression-weighted average of individual gene embeddings.

### CellHermes as a predictor to enable multi-task biological prediction

Traditionally, biological models are designed for specific downstream tasks, each requiring distinct input and output formats. The diversity of tasks and required designs makes it challenging to build a single model capable of handling diverse prediction problems. In this stage, we leverage the power of LLMs to unify heterogeneous downstream tasks into a common natural language framework, enabling CellHermes to function as a predictor across multiple biological domains (Fig. 4a). To evaluate the ability of CellHermes, we propose BioUniBench, a new benchmark integrating 7 databases, including Perturbase^42^, Depmap^43^, BioGRID^4^, CellMarker^68^, CTD^69^, Orphanet^70^, and DisGenNet^71^ (Supplementary Note 5). These resources encompass perturbation effects, fitness landscapes, molecular interactions, cell identity, and disease/phenotype associations—providing a biologically comprehensive and diverse testbed to evaluate CellHermes’s ability to unify heterogeneous downstream tasks within a single predictive framework (Fig. 4b and Supplementary Fig. 13). From these resources, we built 10 training datasets and 13 testing datasets (Supplementary Note 5), with task-specific instructions detailed in the Supplementary Note 6. We used these examples in BioUniBench to illustrate the potential of CellHermes as a universal biological predictor by transforming different biological resources into natural language.

**Fig. 4.**
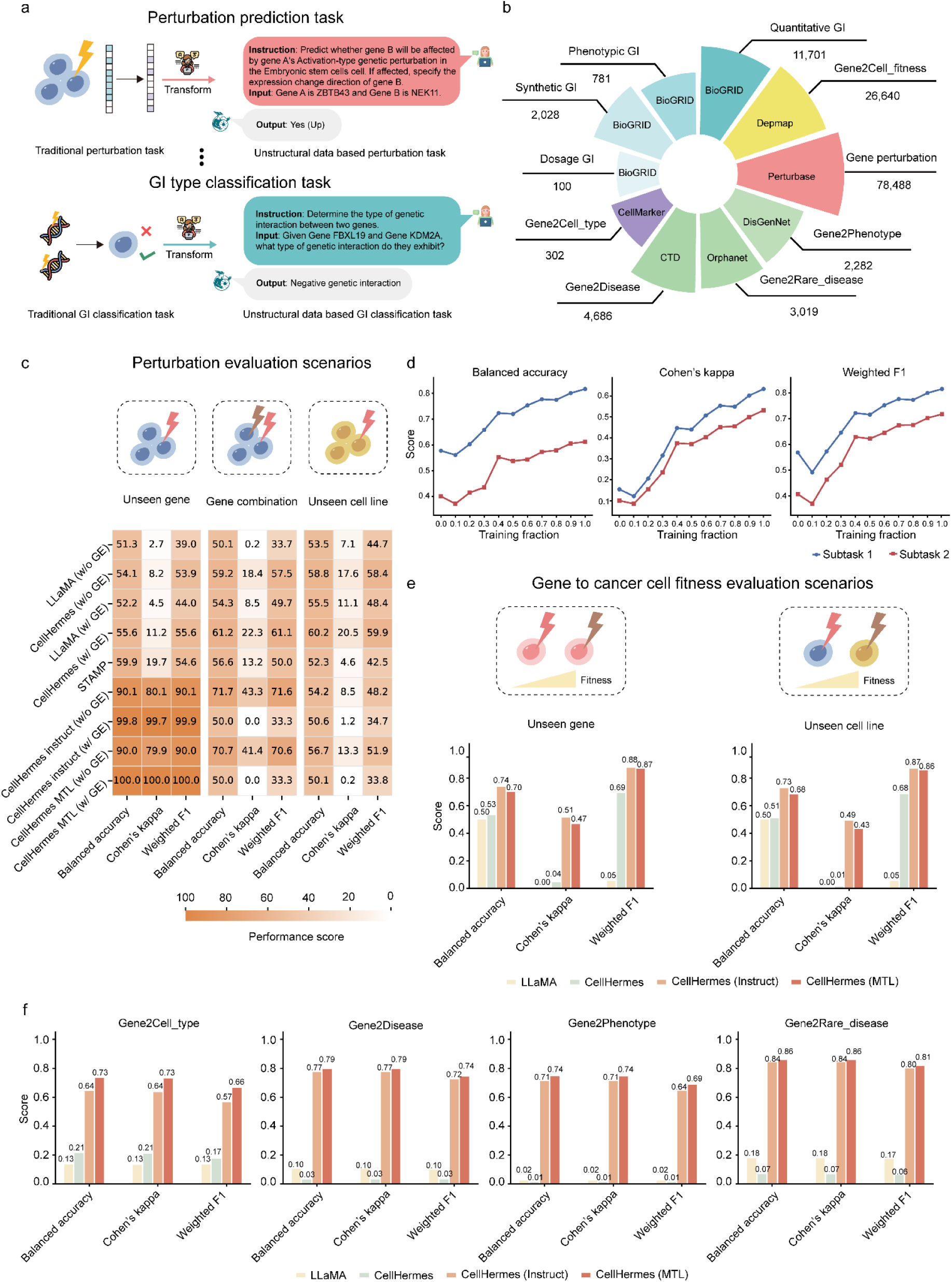
| CellHermes as predictor for multi-task predictions. a. Schematic illustration of transforming different biological downstream prediction tasks into question-answer pairs. b. Composition of multi-task training instruction datasets. Task names are indicated above the horizontal lines, dataset sizes are shown below, and data sources are annotated within the corresponding pie chart segments. c. Performance of different methods on sub-task 1 (DEG prediction) within the genetic perturbation task, evaluated across three scenarios: unseen genes, unseen gene combinations, and unseen cell lines. In this case, we evaluated three variants: CellHermes (zero-shot), CellHermes-instruct (fine-tuned on individual tasks), and CellHermes-MTL (jointly fine-tuned across all tasks). d. Scaling law of CellHermes on genetic perturbation prediction; The x-axis denotes the number of training genetic perturbations. The y-axis represents the model’s prediction performance; color denotes the different sub-tasks. e. Performance of different methods on gene-to-cell fitness prediction tasks across unseen gene and unseen cell line scenarios. f. Performance of different methods on gene set-based classification tasks in BioUniBench, including cell type, disease, phenotype, and rare disease classification.

Quantitative results showed that CellHermes can handle different tasks within one model without compromising performance on individual tasks, while also benefiting from cross-task knowledge transfer. In this setting, CellHermes-instruct denotes model instruction fine-tuned on individual tasks, whereas CellHermes-MTL denotes model jointly fine-tuned across all tasks. For the perturbation prediction task^17,72^, we adopted the structured prediction strategy used in STAMP^73,74^, consisting of differentially expressed gene (DEG) prediction, gene expression change direction prediction. As we employed CellHermes as a language-based predictor, explicit expression values were not incorporated at this stage. We evaluated performance under several challenging settings, including unseen genetic perturbations, unseen gene combinations, and unseen cell lines^17^ (Fig. 4c and Supplementary Fig. 13a). Across all three scenarios, CellHermes in the zero-shot setting outperformed baseline models, regardless of whether the cell’s gene expression profile was included. With instruction fine-tuning, performance further improved within the same cell line, but decreased when generalization across different cell lines. This drop, especially pronounced when expression profiles were included, may reflect overfitting due to the limited diversity of cell line data (Fig. 4c and Supplementary Fig. 13a). Indeed, scaling analysis confirmed that an increasing number of training perturbation samples consistently improved performance, suggesting a clear scaling law^75^ of CellHermes (Fig. 4d and Supplementary Fig. 13a).

We next examined the model performance on cell fitness prediction using DepMap, which includes 1,178 cell lines (Supplementary Note 5). Here, CellHermes with instruction fine-tuning achieved superior performance in both same-cell and cross-cell scenarios (Fig. 4e). The larger diversity of cell lines in this dataset likely reduced overfitting compared to perturbation prediction. Importantly, multi-task fine-tuning did not compromise performance much in this task. Furthermore, we evaluated the model performance across 4 different GI-type tasks. CellHermes with instruction fine-tuning can also substantially improve the model’s performance, and multi-task instruction fine-tuning maintained strong performance without a trade-off (Supplementary Fig. 13b).

Finally, we tested CellHermes on gene set-related downstream tasks, including cell type identification, gene-disease or gene-phenotype associations. In this dataset, CellHermes with instruction fine-tuning can largely improve performance on these tasks, while CellHermes with multi-task instruction fine-tuning can further enhance the model’s performance, benefiting from the knowledge transfer between similar tasks (Fig. 4f). Altogether, these results demonstrate that CellHermes unifies heterogeneous downstream tasks into a single predictive framework by leveraging the comprehensiveness of language modeling, thereby achieving competitive or superior performance, while exploiting cross-task synergies to improve generalization.

### CellHermes as an explainer to provide interpretable explanations for biological discovery

Beyond LLM as encoder and predictor, LLM can also be used as an explainer by leveraging its understanding and reasoning capability. To illustrate this, we applied CellHermes to predict the tumor reactivity of T cells based on their transcriptomic profiles and then interrogated the model’s predictions using both attention-based analysis and text-based reasoning (Fig. 5a and b), as identifying tumor-reactive T cells and uncovering the mechanisms underlying the reactivity is of major biological and clinical importance. We assembled a dataset^76^ of more than 1,000 T cells with experimentally annotated reactivity labels from melanoma patients and constructed instruction-label pairs to fine-tune CellHermes for tumor-reactivity classification (Supplementary Note 7).

**Fig. 5.**
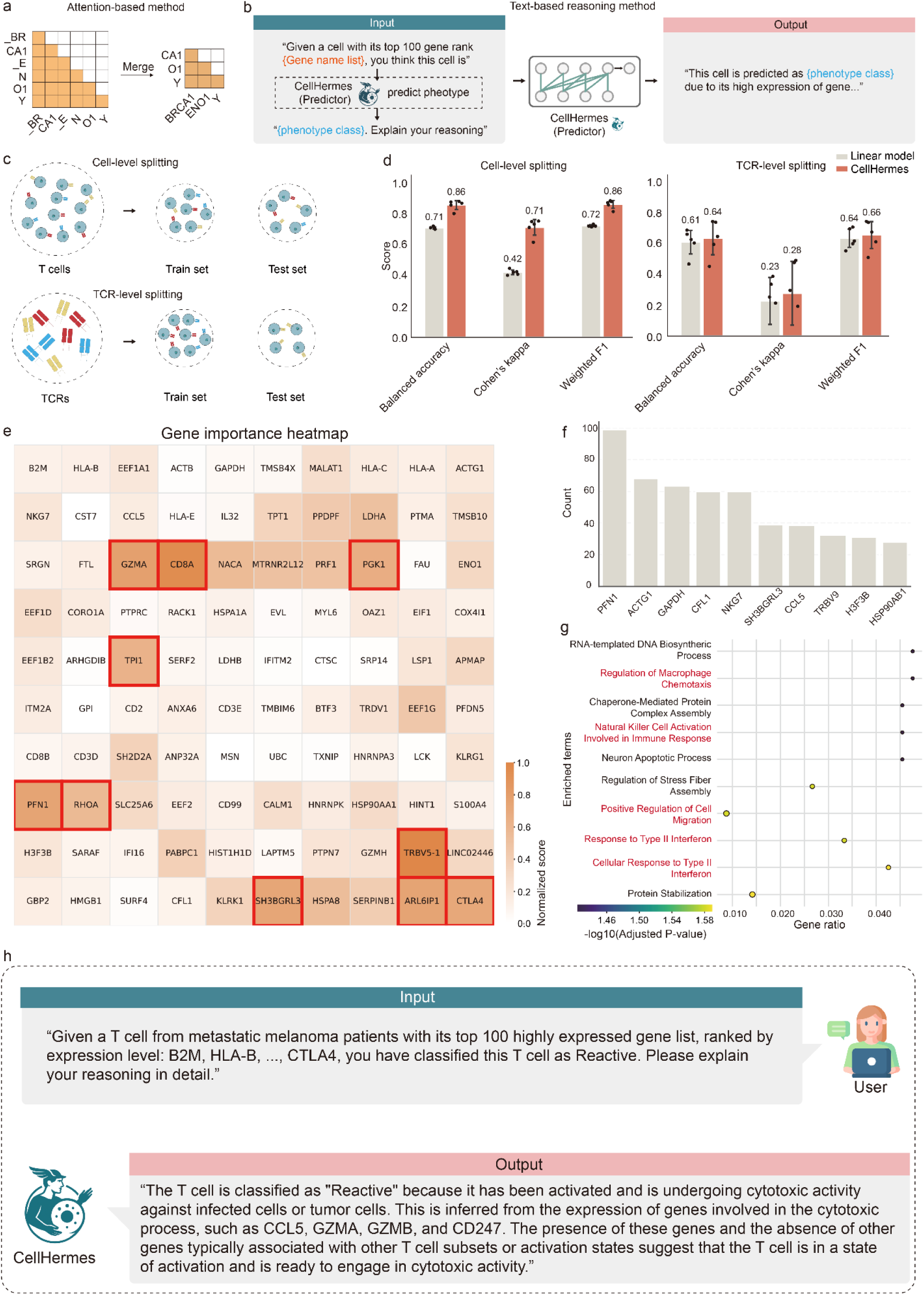
| CellHermes as an explainer for interpreting prediction results. a. Schematic illustration of the attention-based method for interpreting prediction results, where attention matrices from the transformer layer are extracted and sub-tokens merged into gene-level contributions. b. Schematic illustration of a text-based reasoning method for interpreting prediction results, where CellHermes is prompted to generate explanations for its predictions. An example workflow demonstrates how CellHermes is prompted to explain its predictions. First, transcriptomic data is provided as input, and CellHermes predicts the phenotypic class. Then, a prompt is constructed using both the original transcriptome and the predicted phenotype to instruct CellHermes to generate a reasoning-based explanation. In this context, “CellHermes (predictor)” refers to the model that has been instruction fine-tuned specifically for the cell phenotype classification task. c. Schematic illustration of different data splitting strategies for evaluating T cell reactivity prediction (cell-level splitting vs TCR-level splitting). d. Performance of different methods on T cell reactivity prediction under cell-level and TCR-level splits. e. Heatmap of gene importance score for the T cell reactivity, derived from attention weights. f. Frequency of recurrent top-ranking genes by gene importance scores across all cells. g. GO BP enrichment analysis for identified genes from attention weights. These genes are recurrent top-ranking genes across all cells. h. Case analysis for text-based reasoning output illustrating why a T cell was predicted as tumor-reactivity by CellHermes.

To evaluate the predictive performance, we considered two evaluation schemes: (1) cell-level splitting, where cells are randomly divided into training and testing sets, and (2) TCR-level splitting, where cells shared the same TCR are placed into the same partition (Fig. 5c). The latter represents a more challenging scenario, as cells with identical TCRs often share highly similar transcriptomic profiles. In both settings, CellHermes with instruction fine-tuning outperformed a traditional linear model (Fig. 5d). Overall performance was lower in the TCR-level split, likely reflecting the complexity of T cell reactivity, which depends not only on transcriptomic signatures but also on TCR protein structures and other contextual factors.

To explain its predictions, we adopted a dual approach of attention-based analysis and text-based reasoning. We first examined attention weights within the model. Because the LLaMA-3.1-8B-Instruct tokenizer is based on byte-pair encoding (BPE)^77^, gene tokens can be split into multiple pieces (Methods). Therefore, we merged sub-tokens into a single gene-level contribution score. We then focused on the final transformer layer and selected the most informative attention head^78,79^, determined by entropy, to calculate per-gene importance scores (Methods). For an example cell, this analysis revealed a set of genes with high contribution to tumor-reactivity predictions (Fig. 5e). Aggregating across cells, we identified recurrent top-ranking genes, which were enriched for pathways related to T cell activation according to GO BP analysis (Fig. 5f and g). The expression profile further confirmed that these genes were differentially expressed between reactive and non-reactivated T cell populations (Supplementary Fig. 14). Beyond model-internal attention analyses, we also harnessed the reasoning capabilities of CellHermes to generate human-readable explanations. By providing designed prompts that asked the model to justify its predictions, CellHermes was able to give its reasoning process underlying its classification of a given T cell as tumor-reactive (Fig. 5h). This dual approach of integrating architectural attention mechanisms with LLM-based natural language reasoning demonstrates the interpretability of CellHermes and its potential utility as a biological discovery tool.

## Discussion

In this study, we introduce CellHermes, a general-purpose biological language model that unifies heterogeneous data into a single LLM framework. By transforming structural biological information into natural language-like instructions, CellHermes integrates diverse modalities and tasks, functioning as an encoder, predictor, and explainer. Our results demonstrate that, instead of training from scratch, CellHermes achieves competitive performance in representing biological entities with LORA fine-tuning on existing natural language-LLM, supports multi-task biological prediction, and provides interpretable explanations for outputs. These findings highlight the potential of using natural language as a unifying medium for modeling the complexity of omics. As a proof-of-concept framework, CellHermes is readily extensible to additional biological resources and tasks, though in this study, we focused on representative examples for illustration. We note that the representational concept for transcriptomics underlying CellHermes—treating cells as ranked gene ‘sentences’ for use with pretrained LLMs—was originally introduced by Cell2Sentence^7^, itself following the ranked-gene formulation of Geneformer^20^. Our contribution lies in expanding this representation beyond transcriptomics to integrate multimodal omics (including interaction graphs) within a unified, question–answer–based learning framework, implemented through efficient LoRA fine-tuning rather than large-scale autoregressive training.

CellHermes allows for various extensions, for which we found the following most promising. First, its training corpus was relatively modest compared to the scale of available biomedical data. Large and more diverse multimodal corpora, including proteomics, epigenomics, metabolomics, spatial omics, and histology images, will likely further enhance model generalization. Second, CellHermes is presently built on the LLaMA-3.1-8B-Instruct due to computational constraints. Exploring larger open-source language models and systematically benchmarking various existing language models, such as Deepseek^80^ and Qwen^81^, on omics understanding represent an important next step. Third, to further advance open research in this direction, we will release and keep updating BioUniBench by curating and providing more high-quality question-answer pairs for developing and benchmarking text-based biological foundation models. Finally, this study emulates self-supervised learning^82^ to align the omics data with existing LLMs by converting it into text, a method that discards actual numerical values of data. A recent study, CellWhisperer^83^, aligns cellular data with language models using contrastive learning between Geneformer^20^ and BioBERT^84^ to enable rapid cell retrieval and exploration. However, the performance may be constrained by the capability of the single-cell foundation model it used. Exploring other alignment techniques, such as bridging domain-specific models with LLMs^85^, and using reinforcement learning for further improving model’s reasoning ability^86^ could provide a solution for these limitations. Overall, CellHermes defines a modeling paradigm in biology, in which LLMs serve as a connective layer between omics data and reasoning. This may facilitate the construction of omics-specific expert model for agentic assistant and the research loop connecting hypothesis generation with in-lab experimental validation^87^, and provide a potential solution for construction of virtual cells^88^.

## Methods

### Pretraining data collection and preprocessing

For a single-cell transcriptome dataset, we transform it into a gene sequence structure for each cell to enable the use of LLMs. Let ∈ *R*^*n* × *k*^ denote a matrix with *n* rows and *k* columns corresponding to *n* cells and *k* genes, where *C*_*ij*_ represents the number of mRNA molecules counted for gene *j* in cell *i*. Then the count matrix was row-normalized (summing to 10,000) and log-normalized by Scanpy, yielding the preprocessed transcript matrix *C*. For the Tabula Sapiens dataset, which contains 61,759 genes, we filtered out mitochondrial and ribosome genes, as well as genes without gene symbols. We then only retained the protein-encoding genes, resulting in a final set of 19,306 genes. Next, we ranked the gene expression values for each cell and truncated the top 100 genes for each cell *i*, denoted as *S_i_*.

For the Protein-protein interaction dataset, we downloaded the datasets from BioGRID (version:4.4.240). As we focused on the protein-encoding genes in human, we filtered the dataset with parameters {’Organism Name Interactor A’=’Homo sapiens’, ‘Organism Name Interactor B’=’Homo sapiens’, ‘Experimental System Type’=’physical’}, resulting in a final set of 19,977 genes. Then the PPI network can be represented as *G* = (*V*, *A*), where *V* is the set of genes, 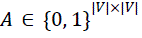 is the adjacency matrix.

### Self-supervised learning (SSL) emulation instruction dataset construction

Structural biological data, including sequence and graph in this study, were converted into natural language descriptions using their specific instruction templates. For sequence data, we designed four templates for masked language modeling (MLM) and autoregressive prediction, where gene identities were serialized into natural language “sentences” based on gene expression levels. In MLM, given a tokenized sequence of genes *S* = (*s*_1_, *s*_2_, …, *S_T_*), a subset of tokens *M* ⊂{1, …, *T*} was replaced with a mask token [*MASK*]. The model was trained to minimize the negative log-likelihood of recovering the masked tokens

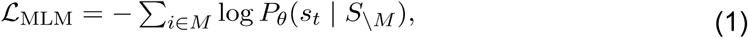

where *P*_θ_ represents the probability distribution defined by the model parameters θ.

In the autoregressive task, the model was trained to predict the next token from its preceding context,

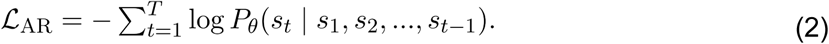

We use natural language to emulate these two objectives by constructing the question-answer pairs (Supplementary information). In MLM, the instruction specifies which tokens are masked, and the answer corresponds to the missing tokens. In AR, the instruction is the prefix sequence, and the answer is the next tokens. Then, by virtue of the flexibility of natural language, we can easily incorporate all meta information *o* of the cell into the objective, obtaining 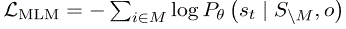 and 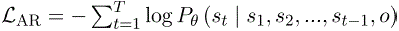. To enhance linguistic diversity, we used ChatGPT-4o to generate 20 paraphrased prompt variants for sequence-level instruction templates. Specifically, for each task, we begin with a human-written description, which then serves as an input to ChatGPT-4o for generating diverse instructions templates, capturing a broad range of natural human question-framing styles.

For graph-based data, we designed five templates for masked node prediction and ten templates for mask link prediction, each varying by hop-level neighborhood information (Supplementary information). In masked node prediction, for a graph *G* = (*V*, *A*) with an adjacency matrix *A*, a node *v* ∈ *V* was masked and predicted from its neighborhood *N_k_*(*v*) with *k*-hops:

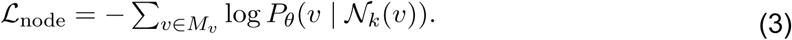

In masked link prediction, an edge (*u*, *v*) ∈ *E* was masked, and the model was trained to predict its presence given the local context:

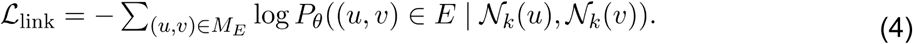

We use natural language to emulate these two objectives by constructing question-answer pairs, considering different scenarios (Supplementary information). In masked node prediction, the instruction encodes the graph context with a masked node, and the answer is the identity of the node. In masked link prediction, the instruction specifies the graph neighborhood with a missing edge, and the answer is the predicted link. Prior studies have demonstrated that incorporating up to three-hop connectivity provides sufficient structural context for efficient learning, whereas higher-order neighborhoods contribute marginal gains and may even introduce noise. Accordingly, the maximum neighborhood depth in our graph templates in this study was set to three.

Altogether, these processes yielded a total of 2,272,436 question-answer pairs for MLM, 2,272,436 for autoregressive modeling, 99,860 for masked node prediction, and 1,455,846 for masked link prediction (Supplementary Note 1). We randomly subsampled different numbers of samples from each dataset to test its impact on model performance. The final corpus, consisting of randomly sampled 0.1 million sequence-based instructions and 0.2 million graph-based instructions, was used to pretrain CellHermes.

### CellHermes model pretraining

By casting each task into a question-answer format, all self-supervised learning (SSL) tasks reduce to predicting a textual answer given a textual instruction. Thus, we can use a universal loss function to jointly learn across heterogeneous modalities and tasks within a single framework. We built CellHermes by instruction fine-tuning a pretrained LLaMA-3.1-8B-Instruct model using LoRA adaptation. Let *x* denote the instruction component and *y* the corresponding answer in the instruction dataset, with a combined sequence length of *L*. The training objective is to minimize the negative log-likelihood:

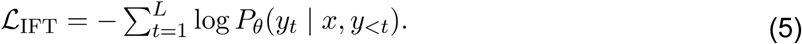

In LoRA, instead of updating the full weight matrix *W* ∈ *R^d×k^*, the update is parameterized as a low-rank decomposition:

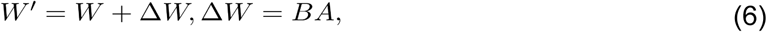

where *A* ∈ *R^r^*^×*k*^ and *B* ∈ *R^d^*^×*r*^ are learnable matrices with rank *r*≪*min*(*d*, *k*). During training, only *A* and *B* are updated while *W* remain frozen. Although LLaMA-3.1-8B-Instruct was used as the backbone in this study, our constructed corpus is compatible with any open-source large language model. Hyperparamers for tuning are listed in Supplementary Note 8.

### CellHermes as an encoder for embedding biological entities

#### Gene embeddings

Because LLMs are inherently generative and not designed as embedding models, we adapted CellHermes into an embedding framework by extracting hidden state representations. The rationale of this approach is that, through the causal attention mechanism, the last token attends to all preceding tokens. Specifically, gene embeddings are extracted by taking the final sub-token embedding from the last transformer layer when the complete gene name (e.g. *BRCA1*) was provided as input. Unlike GenePT, which uses NCBI gene annotations, CellHermes directly encodes gene symbols consistent with the gene format with previous single-cell foundation models. Each gene was embedded into a 4096-dimensional vector space, corresponding to the hidden dimensionality of LLaMA-3.1-8B-Instruct model. Then, these gene embeddings are benchmarked against existing models in the same downstream tasks, including gene property prediction tasks and functional similarity tasks. Gene property prediction tasks comprised classification of long versus short range TFs, dosage sensitivity, bivalent versus non-methylated genes, Lys4-only-methylated versus non-methylated genes and gene-gene interactions. Full detailed dataset descriptions and task details are provided in the Supplementary information. In functional similarity tasks, we evaluated it using two approaches, following prior studies. First, to test whether embeddings captured known functional relationships, we calculated pairwise cosine similarity between all gene embeddings and compared these with semantic similarities derived from curated functional annotation databases, including GO Biological Process (GO BP), GO Cellular Component (GO CC), GO Molecular Function (GO MF), and KEGG pathways. Correlations between embedding-based similarities and semantic similarities were quantified using Pearson’s correlation coefficient. Second, to evaluate the ability of embeddings to recover functional groupings, we applied K-means clustering to the gene embeddings with the number of clusters *K* ranging from 20 to 80. For each configuration, we performed gene set enrichment analysis to assess functional coherence with *gseapy* package. The enrichment cluster percentage was defined as the proportion of clusters significantly enriched (FDR-adjusted p value < 0.05) for one or more terms across the above annotation databases.

#### Cell embeddings

For cell-level representation learning, we considered two strategies adopted by GenePT: (1) Sentence-based embedding (CellHermes-s), which encodes cells by extracting the last sub-token embedding from the final transformer layer of a “cell sentence” (ranked gene names); and (2) weighted gene-embedding average (CellHermes-w), a weighted average of gene embeddings based on expression levels. Performance of both strategies was benchmarked against existing models across five datasets: Aorta, Immune, Lung, Pancreas, and PBMC. Full detailed dataset descriptions are provided in the Supplementary information. Biological preservation and batch correction were quantified using *scib-metrics*.

#### Cell-specific gene network construction

To further assess the utility of CellHermes representations, we constructed cell-specific gene networks. For each cell, pairwise similarity between gene embeddings was computed and a KNN graph was then generated, where each gene was connected to its *k*=15 most similar neighbors. This procedure was implemented by *sc.pp.neighbors* in *Scanpy.* By comparing networks across different cell types, we first calculated the node degrees of each gene and identified those showing the greatest degree change between cell types. To investigate whether constructed networks could capture cell-type-specific gene modules, we performed community detection and module similarity analysis. For each cell type, Louvain clustering was applied to the corresponding cell type-specific networks to identify gene modules. The similarity between modules from different cell types was quantified using the Jaccard Index, defined as:

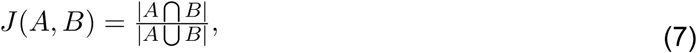

where A and B are two gene modules. In this study, we considered modules to be cell type-specific if their Jaccard Index was less than 0.4 with all modules of the other cell type. Then, we used *sc.tl.score_genes* in *Scanpy to* give an overall expression value for each gene module and identified the differentially expressed gene modules by using *sc.tl.rank_genes_groups* with the Wilcoxon test. Functional coherence of these modules was evaluated using GO BP enrichment with the *gseapy* package. For comparison, we also constructed co-expression networks by computing the Pearson correlation coefficient between every gene pair and connecting each gene to its *k*=15 most correlated neighbors using the same KNN procedure.

### CellHermes as a predictor for multi-task prediction

Seven databases were collected for multi-task evaluation: Perturbase, DepMap, BioGRID, CellMaker, CTD, Orphanet, and DisGeNET. These encompassed genetic perturbation prediction, cell fitness prediction, genetic interaction classification, cell type identification, and gene-disease/phenotype association. After preprocessing, we constructed 10 training datasets and 13 testing datasets. Task-specific instructions were generated for each dataset (Supplementary information).

For the perturbation prediction task, we adopted the STAMP framework, evaluating differential expression DEG prediction and expression change direction prediction. Additional testing scenarios included unseen genes, unseen gene combinations, and unseen cell lines. In this task, we benchmarked multiple model configurations:

(1) LLaMA (baseline pretrained model, without fine-tuning or gene expression input),
(2) STAMP (baseline linear model),
(3) CellHermes (w/o GE) without fine-tuning and without gene expression profile,
(4) CellHermes (w/ GE)without fine-tuning but incorporating gene expression profile,
(5) CellHermes (instruct w/o GE) with instruction fine-tuning but without gene expression profile,
(6) CellHermes instruct (w/ GE) with instruction fine-tuning and gene expression profile,
(7) CellHermes MTL (w/o GE) with multi-task fine-tuning but without gene expression profile,
(8) CellHermes MTL (w/ GE) with both multi-task fine-tuning and gene expression profile.

For cell fitness prediction, we used the DepMap-derived dataset and constructed two testing scenarios: an unseen gene scenario and an unseen cell line scenario. For genetic interaction classification, we evaluated performance across four different genetic interaction (GI) types. For gene set–based tasks (cell type, disease, phenotype, and rare disease), the objective was to classify gene sets according to their associated category. All of these heterogeneous biological knowledge sources, despite differing input formats, were converted into a unified instruction–response format. The resulting instruction datasets were then used to perform multi-task fine-tuning of CellHermes following Equation (5).

### CellHermes as an explainer for interpretability analysis

To explain predictions, we use both attention-based and text-based reasoning approaches. For attention-based analysis, we extracted attention weights from the final transformer layer for a cell input sequence of tokens *S* = (*s*_1_, *s*_2_, …, *S_T_*). The attention mechanism is defined as:

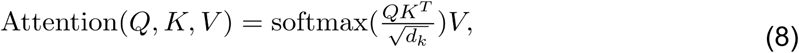

where *Q*, *K*, *V* are the query, key, and value matrices, respectively, and *d_k_* is the key dimensionality. Among all heads, we selected the one with the lowest mean entropy, as this head is typically the most focused and interpretable. We focused on the head with the lowest mean entropy. The entropy for a target position *t* in the head ℎ was computed as:

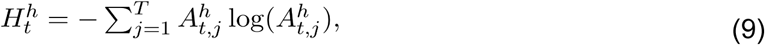

where 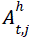 is the attention weight assigned to the source token *j* by the head ℎ for the target token *t*. Because the tokenizer is based on byte-pair encoding (BPE), each gene name may be split into multiple sub-tokens. To obtain gene-level contributions, we aggregated sub-token attention weights into a single per-gene contribution score:

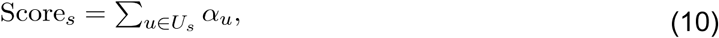

where *U*_*s*_ is the set of sub-tokens for gene *s* and α_*u*_ the attention weight. These scores were then used to rank genes contributing to the model’s prediction and identify genes most responsible for driving predictions. For text-based reasoning, we designed natural language prompts asking CellHermes to justify predictions. The model’s responses were compared with gene-level attention results, providing complementary interpretability from architectural and linguistic perspectives.

## Data availability

All datasets utilised by CellHermes for training, testing, and application examples were obtained from publicly accessible databases. Specifically, the transcriptome dataset of pretraining is available at [https://cellxgene.cziscience.com/collections/e5f58829-1a66-40b5-a624-9046778e74f5], while the PPI graph dataset is available at [https://thebiogrid.org/]. The gene-level downstream tasks data can be access from [https://huggingface.co/datasets/ctheodoris/Genecorpus-30M/tree/main/example_input_files/gene_classification] and the gene-gene interaction network datasets are available at [https://github.com/jingcheng-du/Gene2vec/tree/master/predictionData]. The Aorta^66^ dataset is available at [https://drive.google.com/drive/folders/1LgFvJqWNq9BqHbuxB2tYf62kXs9KqL4t?usp=share_link], Immune^61^, Lung^61^, Pancreas^61^ datasets are available at [https://doi.org/10.6084/m9.figshare.12420968] and PBMC^67^ dataset is available at [https://ngs.sanger.ac.uk/production/teichmann/BBKNN/PBMC.merged.h5ad]. The T cell reactivity dataset^76^ is available in the Gene Expression Omnibus (GEO) under accession number GSE222448. Source data are provided in this paper.

## Code availability

CellHermes is publicly available on GitHub [https://github.com/theislab/CellHermes] and model weights can be obtained on [https://huggingface.co/collections/tongjideltalab/cellhermes], under the GPL-3.0 license, together with usage documentation.

## Acknowledgments

Q.L. was supported by the National Key Research and Development Program of China (Grant No. 2025YFC3409300), the National Natural Science Foundation of China (Grant No. T2425019, 32341008), Shanghai Pilot Program for Basic Research, Shanghai Science and Technology Innovation Action Plan-Key Specialization in Computational Biology, Shanghai Shuguang Scholars Project, Shanghai Excellent Academic Leader Project, Shanghai Municipal Science and Technology Major Project (Grant No. 2021SHZDZX0100) and Fundamental Research Funds for the Central Universities. Y.G. was supported by National Natural Science Foundation of China (Grant No. 324B2013). F.J.T. is supported by the European Union (ERC, DeepCell – 101054957). T.R. is supported by the Helmholtz Association through the Munich School for Data Science. T.R. also receives funding from the Munich Center for Machine Learning (MCML; BMBF; grant #01IS18053A). W.W. is supported by funding from the European Union’s Horizon 2022 HORIZON research and innovation programme (Grant No.101057775).

## Author contributions

Y.G., Q.L. and F.J.T. designed the framework of this work. Y.G. and C.S. provided technical support. Y.G., W.W., D.K., W.Z. and Z.L. explored downstream biological applications. Y.G., W.W., Y.Z. and D.K. performed the analyses. Y.G., Q.L., W.W and T. R. wrote the manuscript with the help of other authors. All authors read and approved the final manuscript.

## Competing interests

The authors declare the following financial interests/personal relationships which may be considered as potential competing interests: F.J.T. consults for Immunai, CytoReason, BioTuring, Genbio and Valinor Industries, and has ownership interest in RN.AI Therapeutics, Dermagnostix, and Cellarity. The remaining authors declare no competing interests.

## Notes

### Summary of Updates

Revised Figures and made some statements more clearer

